# Age and sex influence seed dispersal of native and non-native plants by Lion-tailed Macaques *Macaca silenus*

**DOI:** 10.1101/2024.12.09.627456

**Authors:** K Bindu, Honnavalli N. Kumara, Rohit Naniwadekar

**Author notes:** Corresponding authors:Bindu K. and Rohit Naniwadekar, Address: Nature Conservation Foundation, 1311, “Amritha”, 12th Main, Vijayanagar 1st Stage, Mysuru, Karnataka 570017, India.

## Abstract

While interspecific variation in seed dispersal, a critical ecosystem process in tropical forests, is relatively well-studied, intraspecific variation as a consequence of differences in body size, foraging behaviours, and ranging patterns among age-sex categories within a species is relatively understudied. Among vertebrates, primates play a critical role in seed dispersal and exhibit behavioural differences between age and sex categories, making them a suitable study system for intraspecific variation in seed dispersal. Lion-tailed macaques *Macaca silenus*, an endemic and predominantly frugivorous primate species in the Western Ghats Biodiversity Hotspot, provide an excellent model for such studies. We examined the influence of age and sex on 1) the diversity and 2) the quantity of native and non-native fruits consumed, 3) the number of seeds dispersed, 4) seed dispersal distance, and 5) seed deposition substrates in lion-tailed macaques. We conducted over 375 hours of focal animal watches, distributed evenly across adult males, females, and subadults. Our findings showed that subadults consumed a higher diversity of native and non-native fruits than females and males. They dispersed fewer *Ficus* seeds than females. We found differences in the proportions of non-native fruits in the diets of different age-sex categories. Males consumed more *Coffea liberica*, whereas females and subadults fed on *Coffea* and *Lantana*. We found weak evidence suggesting that males were more likely to disperse *Ficus* seeds on trees, which are suitable substrates for *Ficus* establishment. Our study highlights that age and sex significantly influence seed dispersal patterns of native and non-native species by an endemic, frugivorous primate species with potential influence on recruitment.

## 1. INTRODUCTION

Seed dispersal is a key ecological process influencing plant diversity at local, regional, and continental scales (Levine & Murrell, 2003; Plue & Cousins, 2018). In the tropics, animals play a critical role in seed dispersal, dispersing seeds of up to 90% of the plant species (Farwig & Berens, 2012; Howe & Smallwood, 1982). While the variation in seed dispersal outcomes due to differences in the morphology and ecology of different seed disperser species is well-documented, the influence of individual animals’ personalities and behaviour on seed dispersal outcomes is less explored. Intraspecific variation in seed dispersal is relatively understudied in the literature compared to inter-specific variation (Zwolak & Sih, 2020).

Variations in seed dispersal effectiveness within a species may be driven by factors such as size, age, sex classes, social status, or individual personalities (Zwolak & Sih, 2020). These variations can lead to differences in seed dispersal effectiveness, including the number of seeds dispersed, dispersal distance, and the quality of the seed deposition site (Snell et al., 2019). For example, subordinate females in Japanese macaques *Macaca fuscata* may swallow fruits likely to avoid harassment by dominant individuals, thereby providing better seed dispersal services (Tsuji, Campos-Arceiz, et al., 2020; Tsuji & Takatsuki, 2012), highlighting the role of dominance in influencing seed dispersal. In Howler monkeys *Alouatta pigra*, age and sex impact the time spent foraging and the diversity of plant species in their diet, with adults foraging longer and consuming a greater diversity of plant species (Benitez-Malvido et al., 2016). Given the potential influence of age and sex on frugivory and seed dispersal, a recent review emphasised the importance of understanding the linkages between behavioural types and their impact on seed dispersal (Zwolak & Sih, 2020).

Seed dispersal effectiveness encompasses both qualitative and quantitative components (Schupp, 1993; Schupp et al., 2010). The qualitative components include factors like seed handling behaviour, gut treatment of seeds, and the suitability of seed deposition sites, while the quantitative component refers to the number of seeds dispersed. In the context of intraspecific variation, the quantitative component of seed dispersal can be influenced by differences in relative abundance, dietary differences, and body sizes. Meanwhile, the quality component of seed dispersal can be impacted by variations in ranging patterns, physiology and fruit handling among individuals within a species. Age-sex categories within a species often exhibit differences in relative abundance, body sizes, dietary preferences, ranging patterns, and fruit handling (Benitez-Malvido et al., 2016; Celebias et al., 2024; Dhawale et al., 2020). Most studies on intraspecific variation in seed dispersers tend to focus on particular stages of seed dispersal process, e.g., Camargo et. al. 2011 studied variations in the quantity of seeds dispersed by male and female agile opossum *Gracilinanus agilis*. Studies that examine multiple quantitative and qualitative components of the seed dispersal process are relatively limited (but see Tsuji et al.

2020).

Primates are one of the important seed dispersers in tropical forests, dispersing seeds by spitting or through their faeces (Chaves et al., 2011; Sengupta et al., 2020). In addition to age-related differences, primates often exhibit sexual dimorphism, with males typically being larger than females (Plavcan, 2001). Most frugivorous primates are group-living and have a strong social organisation within their troops (Snaith & Chapman, 2007). The combination of dominance hierarchies and morphological differences among individuals in frugivorous primates makes them an ideal study system to examine the intraspecific variations in seed dispersal patterns. Intraspecific variations in primate behaviour and social status directly influence seed dispersal patterns (Tsuji, Campos-Arceiz, et al., 2020).

The lion-tailed macaque *Macaca silenus* (LTM) is an endemic, habitat specialist, primarily arboreal, and frugivorous primate found in the central and southern portions of the Western Ghats Biodiversity Hotspot in India (Singh, Singh, et al., 2002; Umapathy & Kumar, 2000). Fruits are a significant component of LTM’s diet, constituting up to 70% of consumed food items (Santhosh et al., 2015; Singh, Kumara, et al., 2002). Their diet includes native fruit species such as *Artocarpus heterophyllus*, *Cullenia exarillata*, *Ficus* spp., *Litsea floribunda*, and *Syzygium cumini*, among others. In the severely fragmented rainforests of Anamalai hills, LTMs also consume exotic species, including *Persea americana*, *Maesopsis eminii*, *Coffea canephora*, *Coffea liberica*, *Lantana camara*, and *Psidium guajava* (Dhawale et al., 2020; Menon & Poirier, 1996; Singh, Kumara, et al., 2002). LTMs exhibit a social system characterised by despotism and a linear dominance hierarchy (Singh et al., 2006, 2011). An aggressively structured dominance hierarchy exists among males, females, and between the sexes (Singh et al., 2006, 2011, 2011).

Furthermore, male and female LTMs have been reported to exhibit different levels and types of curiosity, with females demonstrating a greater propensity to locate food resources (Rouff et al., 2005). Given the importance of fruits in their diets and their distinct social hierarchy, LTMs are a suitable species for studying intraspecific variation in seed dispersal.

Puduthottam is a private, degraded rainforest fragment that has the highest density (>100 individuals/km^2^) of LTMs among the 45 rainforest fragments in the Anamalai hills of Tamil Nadu (Dhawale & Sinha, 2023; Singh et al., 2002). The LTM troops in this fragment have been extensively studied for several decades and are well-habituated (Dhawale et al., 2020; Singh, Singh, et al., 2002; Umapathy & Kumar, 2000). Additionally, this severely degraded rainforest fragment has a high density of invasive species, constituting a significant part of the LTM diet. This allows us to examine the influence of age and sex on the differential seed dispersal of alien, invasive plant species in a degraded landscape.

Given the intraspecific differences in LTM behaviour, we examined whether different seed dispersal parameters differed across age-sex categories. Specifically, we examined: 1) the diversity of native and non-native fruits foraged by different age-sex categories of LTMs, 2) the relative contributions of native and non-native fruits in the diets of different age-sex categories, the daily distance travelled by individuals of different age-sex categories, 4) the number of *Ficus* seeds deposited by individuals belonging to different age-sex categories, and 5) the substrates where *Ficus* seeds are deposited by different age-sex categories. This study provides valuable insights into the role of the endemic, forest specialist LTM as a disperser of native and non-native plant species, an area that has been underexplored in the literature.

## 2. METHODS

### 2.1. Study area

The lion-tailed macaque is distributed across eight locations in India’s Karnataka, Kerala and Tamil Nadu states (Kurup & Kumar, 1993; Singh, Singh, et al., 2002). The Anamalai Hills of Tamil Nadu harbours the largest metapopulation within LTMs distributional range (Kumara et al. 2014; Singh, 2019). The primary vegetation type of the area is tropical rainforest, occurring at elevations between 600 and 1500 m. Once a contiguous stretch of tropical rainforest, the area has experienced forest loss and severe degradation over the past two centuries due to land clearing

for tea, coffee, and cardamom cultivation. At the centre of these hills lies the Valparai plateau, a 220 km^2^ heterogeneous landscape dominated by tea and coffee plantations interspersed with several rainforest fragments ranging from <10 ha to >100 ha in size (Umapathy & Kumar, 2000).

We conducted the study in one of the larger fragments, Puduthottam, which spans 92 ha. The fragment is bordered by coffee and tea plantations and human settlements, including plantation labour lines and the nearby town of Valparai. Additionally, the Pollachi-Valparai National Highway cuts through the fragment. The macaques use all available habitat types (Dhawale et al., 2020). Puduthottam harbours five LTM troops with approximately 200 individuals (Dhawale & Sinha, 2023). Of these, BK observed individuals from four troops: BT, NTT, RT, and PAP, consisting of ca. 116, 39, 25 and 12 individuals, respectively (Dhawale & Sinha, 2023; Bindu K, Pers. Obs.).

### 2.2. Field methods

The study involved non-invasive behavioural observations of LTMs. We selected the first troop encountered in the field for observation. We restricted observations to individuals from three age-sex categories: adult males, adult females, and subadult males (Table S1, adapted from Singh et al., 2002). We used focal animal sampling methodology, observing a single individual for up to six hours per day. We started the observations from 0700 hr. On days when the troop was not immediately located, focal animal sampling started after we located the troop and a focal individual was identified. We consecutively sampled adult males, females and subadults to ensure interspersion and minimise potential biases arising from temporal variation in foraging across age-sex categories. We sampled an average of 19.8 (SE: ± 1.84) days per month for four months. The total sampling effort encompassed 375.9 hours of focal observations, distributed among the different age-sex categories: adult males, adult females, and subadult males (Table 1). We followed the troop in fragments, plantations, and plantation labour lines. However, we stopped observations when the troop moved into the adjacent town, as they rarely consumed fruits and primarily fed on human-origin food. While our sampling effort varied across troops, it was similar across age-sex categories (Table 1). While our sampling effort was restricted to a single dry season in our study area, our total effort exceeded that of a previous study on intraspecific variation in Japanese Macaques (Tsuji, Campos-Arceiz, et al., 2020). We did not observe subadult females due to difficulty in distinguishing them from adult females and juvenile males.

**Table 1.** Sampling effort (hours) across four troops distributed across three age-sex categories. “NA” indicates no observations for the particular age-sex category within the respective troop.

| Troop ID | Adult Female | Adult Male | Subadult | Total duration (hr) |
| --- | --- | --- | --- | --- |
| BT | 66.50 | 70.71 | 85.57 | 222.8 |
| NTT | 40.41 | 27.51 | 29.67 | 97.6 |
| PAP | 14.57 | 28.10 | 7.40 | 50.1 |
| RT | NA | 4.41 | 0.96 | 5.4 |
|  | 121.48 | 130.72 | 123.6 | 375.9 |

We categorised the observed activities into moving, resting, grooming, vocalisation, foraging, and feeding (modified from Tsuji et al., 2020). When the focal individual engaged in feeding behaviour, we classified the food items into plant, invertebrate, human-origin food, or unidentified material. When feeding on fruits, we collected additional details like the fruit species identity, the number of fruits consumed, and the state of fruit ripeness (ripe/unripe). Ripeness was visually determined based on the fruit colour.

To determine the daily distance covered by focal individuals, we tracked their movements using a Garmin eTrex 20x GPS. To determine the number of seeds dispersed by LTMs of different plant species, we investigated faeces from individuals of known age-sex categories upon observing defecation events. We recorded the substrate of the deposition sites. We followed established methods (Tsuji, Campos-Arceiz, et al., 2020) and identified seeds to the species level, except for *Ficus* and *Coffea* seeds, which require molecular techniques for identification. We analysed only *Ficus* seeds since other seeds appeared infrequently in the faeces.

### 2.3. Analysis

All statistical analysis was performed in R (ver. 4.2.1) and visualised using ggplot2 (Wickham et al., 2007; R Core Team, 2022).

We evaluated sampling completeness for each age-sex category using sample coverage, which estimates the proportion of the total community represented in the sample (Roswell et al., 2021). The sample coverage for all three cohorts (adult male, adult female and subadults) was >99%, indicating sampling adequacy.

To compare the diversity of fruits consumed by the three age-sex categories, we computed Hill-Shannon diversity for native and non-native fruits consumed by each age-sex category using the R package ‘iNEXT’ (T. C. Hsieh, K. H. Ma and Anne Chao, 2015). We bootstrapped the data 50 times to obtain 95% confidence intervals. We compared the diversity metric across three age-sex categories and inferred statistically significant differences if the confidence intervals did not overlap (Cumming et al., 2007). We chose Hill-Shannon (q = 1) over species richness (q = 0) as it incorporates abundance information, unlike species richness, which treats all species equally regardless of their representation in the diet. Since the study focuses on frugivory and seed dispersal, relative abundance information is crucial, as macaques are likely to contribute less to the dispersal of infrequently consumed species.

We performed the Chi-squared test to examine: 1) differences in the proportions of ripe and unripe native and non-native fruits consumed across age-sex categories, and 2) differences in the proportions of different non-native species consumed across age-sex categories.

We used general linear models to determine whether daily movement (per hour) differed across the age-sex categories. The data were log-transformed to approximate normality. We used generalised linear models to examine whether the number of *Ficus* seeds dispersed differed across age-sex categories. Since the data was overdispersed, we fitted a negative binomial regression model. We performed a Chi-squared test to examine the association between age-sex cohorts and the different kinds of substrates (forest floor and tree) where the faeces were found.

## 3. RESULTS

### 3.1. Fruit Diversity in Diet

We recorded 24 species of plants in the diet of LTM (Table S2). The species richness of fruits consumed by females, males, and subadults was 17, 17, and 19, respectively. The Hill-Shannon diversity of fruits in the diets of females, males, and subadults was 6.4, 3.9, and 8.2 species, respectively. Hill-Shannon Diversity differed significantly among the three age-sex categories as inferred from non-overlapping 95% confidence intervals (CI) (Fig. 1a). A similar pattern was observed for native fruit diversity, with subadults feeding the most diverse array of fruits (6.3 species), followed by females (4.2 species) and males (2.8 species). The diversity of non-native fruits consumed by males (3.1 species) was lower than that consumed by females (3.9 species) and subadults (3.7 species), as inferred from non-overlapping 95% CI.

**Figure 1.**
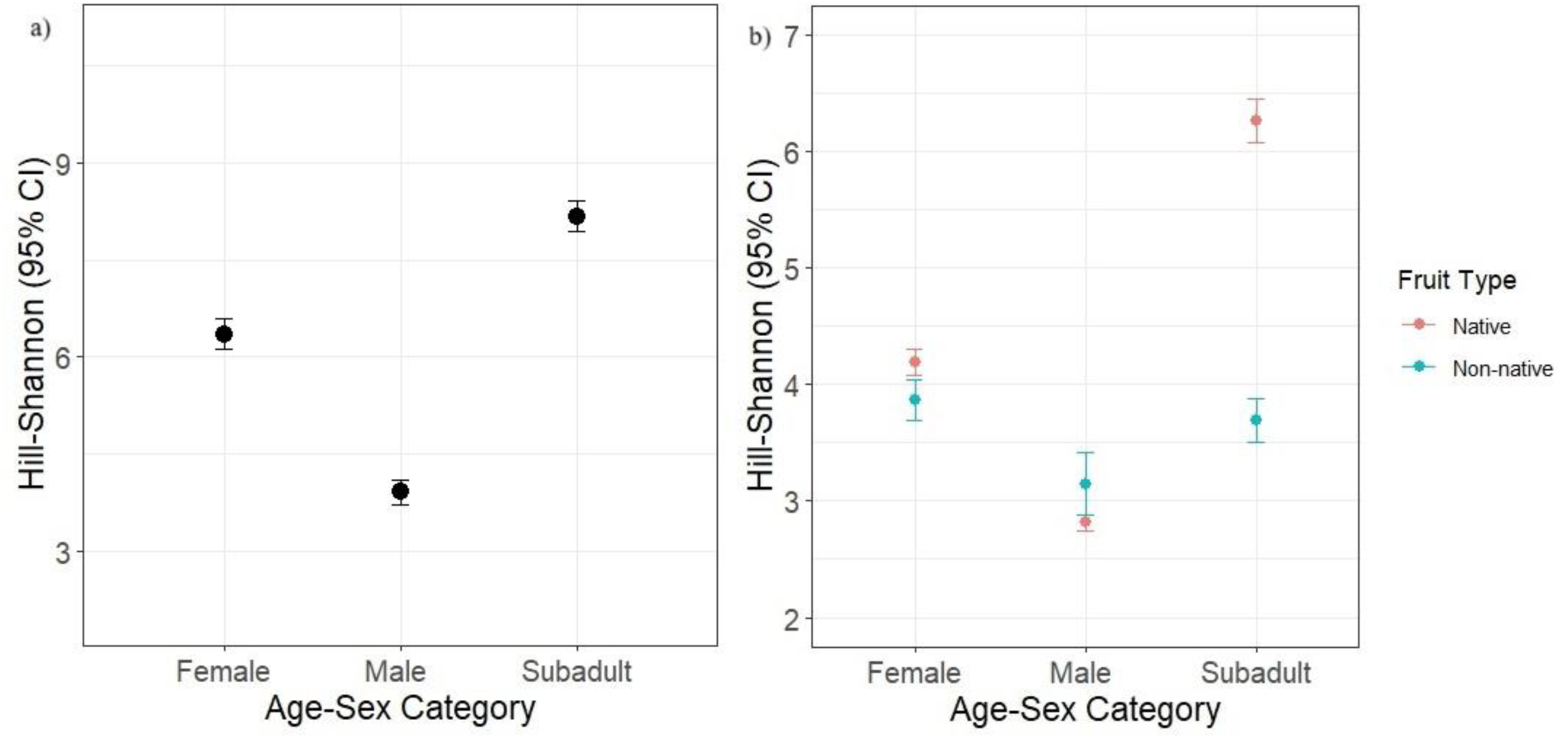
Taxonomic diversity (Hill-Shannon Diversity Measure) of a) all fruits (pooled native and non-native) and b) native and non-native fruits consumed across age-sex categories. Bars represent bootstrapped 95% confidence intervals.

### 3.2. Quantity of Fruits in Diet

We found that subadults consumed a higher proportion of native fruits, while females consumed a higher proportion of non-native fruits than the other age-sex categories (*ꭓ^2^* = 306.31, df = 2, *p* < 0.001; Fig. 2a). Up to 25% of the diet of the female and male LTMs consisted of non-native species. Males consumed a higher proportion of *Coffea liberica* and a lower proportion of *Lantana camara* than females and subadults (*ꭓ^2^* = 425.11, df = 14, *p* < 0.001; Fig. 2b). These two alien species collectively contributed to up to 90% of the invasive in the diet of LTMs.

**Figure 2.**
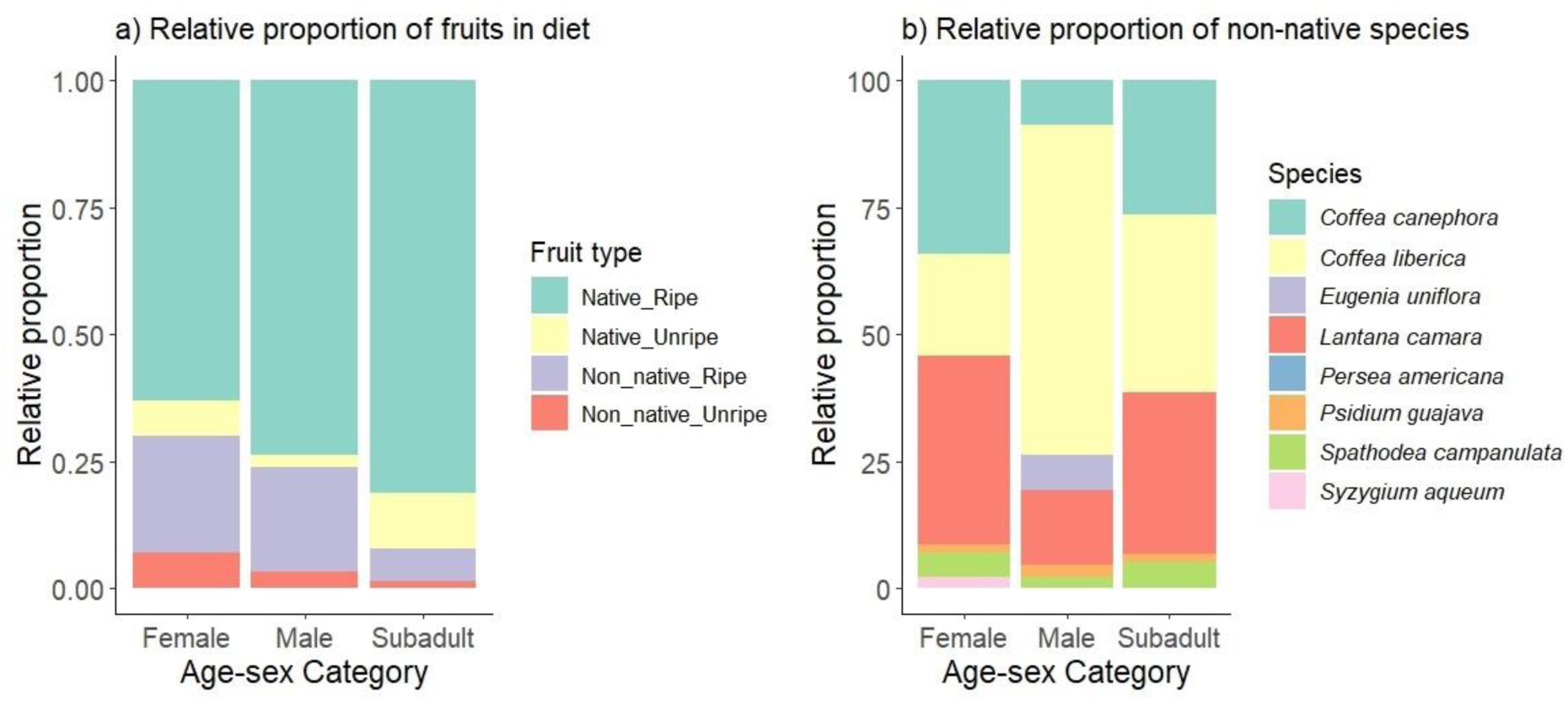
a) Relative proportions of fruits in the diet, showing a lower representation of unripe fruits and higher proportions of non-native and native fruits in the diets of females and subadults, respectively. b) Relative proportions of different non-native species in the diet. Adult males consumed more *Coffea liberica* compared to other alien fruit species, while *Lantana camara* formed a significant part of the diet of females and subadults.

### 3.3. Daily Movement

We recorded 107 tracks of varying lengths during the study. There was no significant effect of age-sex category on daily movement (*F2, 104* = 0.8787, *p* = 0.42; Table S3; Fig. 3).

**Figure 3.**
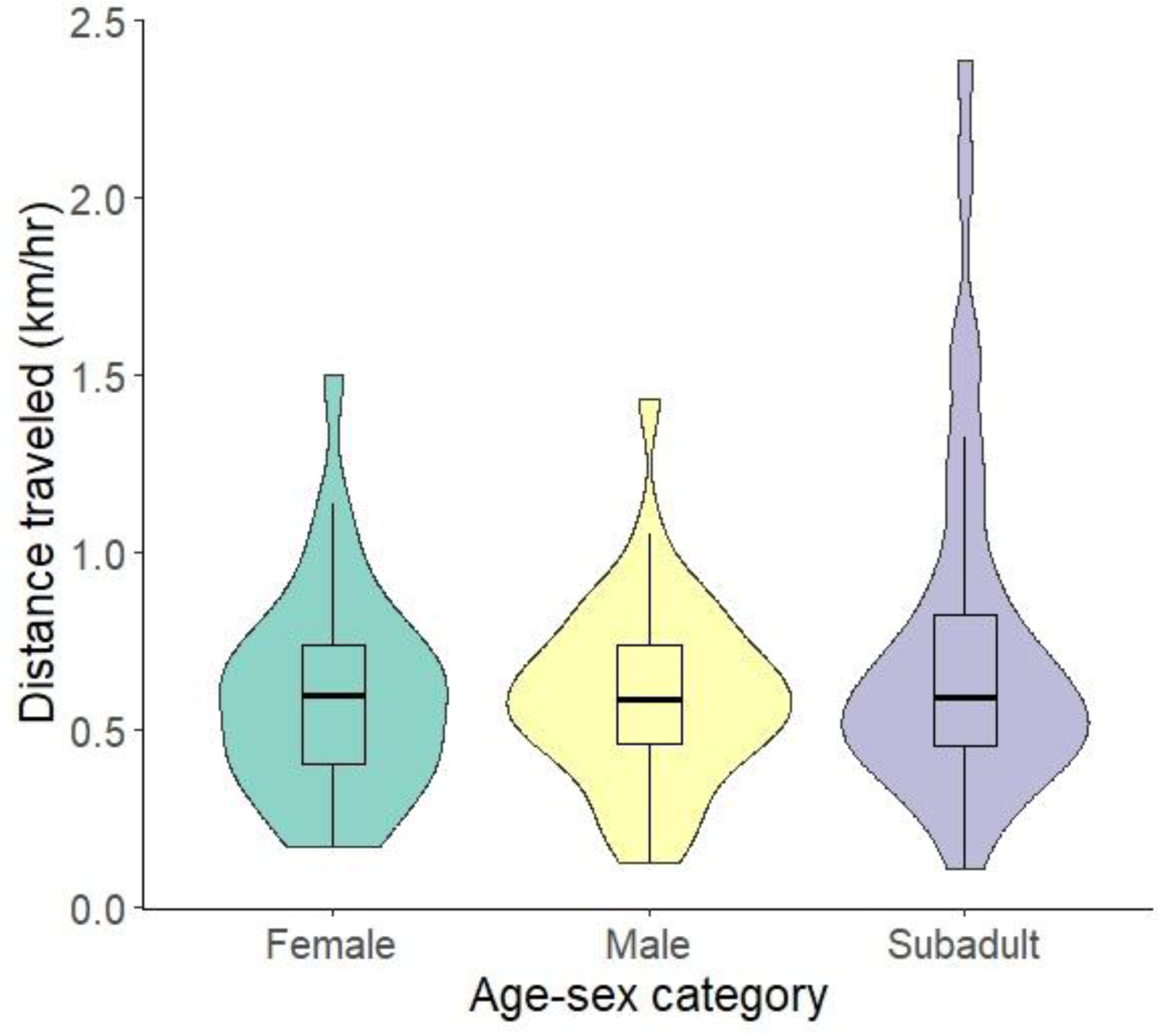
Daily distance travelled (km hr^-1^) across age-sex categories.

### 3.4. Seed Dispersal Quantity

During the study, from 51 fresh faecal deposits investigated, 121,375 seeds belonging to six plant species were identified, including *Ficus* spp., *Coffea* spp., *Maesopsis* e*minii*, *Spathodea campanulata*, *Scurrula parasitica*, and *Lantana camara*. Since seeds from other plant species appeared infrequently in the faeces, only *Ficus* seeds were considered for further analysis.

Subadults dispersed significantly fewer *Ficus* seeds than the other age-sex categories (Fig. 4a, Table S4).

**Figure 4.**
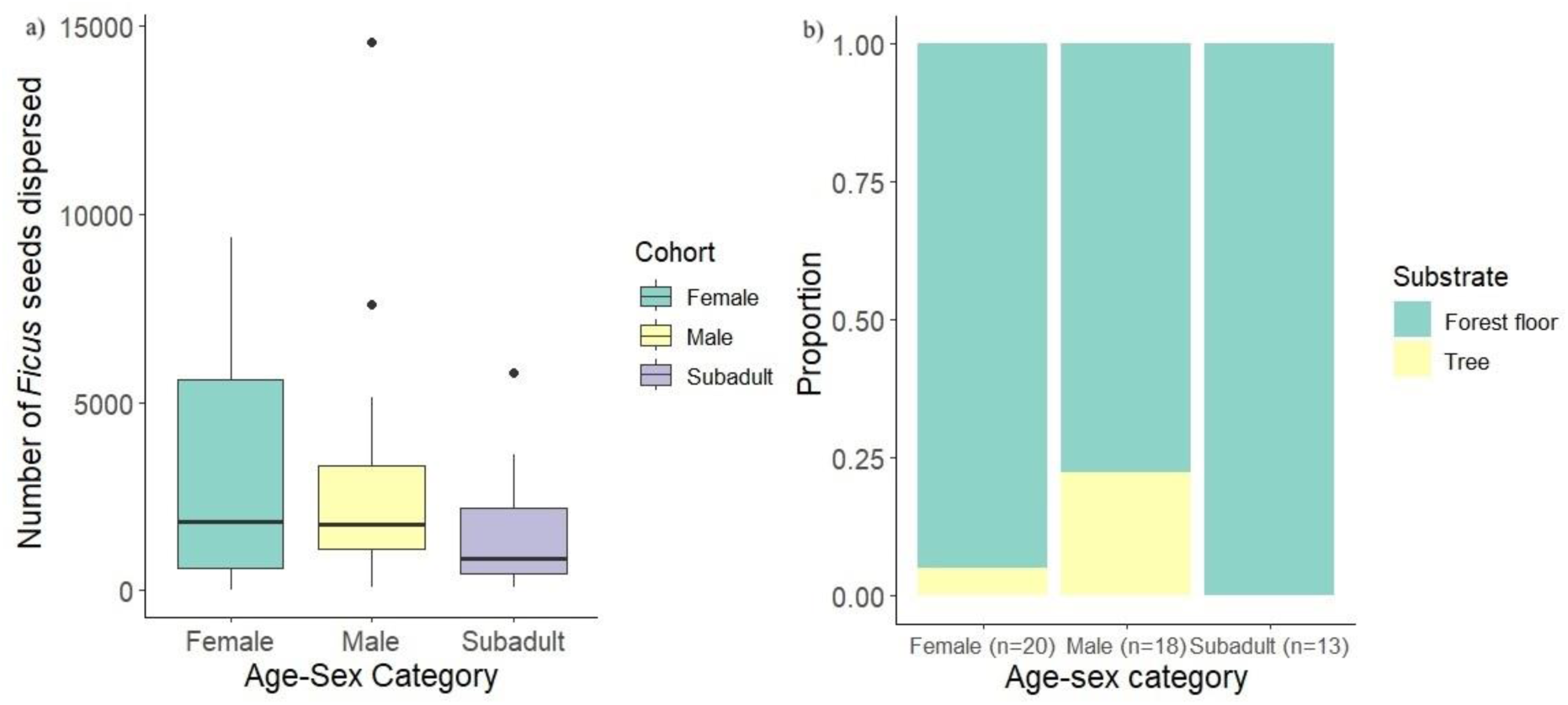
a) Number of *Ficus* seeds dispersed per scat by each age-sex category. b) Relative proportion of seed deposition substrates across age-sex categories.

### 3.5. Seed Deposition Site

We found 51 faecal deposits (females: 20, males: 18, subadults: 13). The relative proportions of faecal deposits across different substrates differed marginally among age-sex categories (*ꭓ^2^* = 11.63, df = 6, *p* = 0.07; Fig. 4b). There was weak evidence that male LTMs were more likely to disperse *Ficus* seeds on trees, which are more suitable habitats for the germination of *Ficus* seeds.

## 4. DISCUSSION

This study is one of the few to examine intraspecific variation in frugivory and seed dispersal by an endemic, forest specialist, frugivorous primate. Our results demonstrate that the age and sex of lion-tailed macaques influence their role in seed dispersal. We found that subadults had higher diversity and greater representation of native fruits in their diet than adult females and males, although they dispersed fewer *Ficus* seeds. In contrast, males tended to disperse *Ficus* seeds on trees, which are more suitable sites for seedling establishment (Laman, 1995; Shanahan, 2016). There were also differences in consumption of non-native fruits, with a greater diversity in the diet of subadults. Males consumed a higher proportion of *Coffea liberica*, whereas females and subadults fed on both *Coffea* and *Lantana*. The variation in the relative abundance of females, males, and subadults within a troop can significantly influence the dispersal patterns of native and non-native seeds across the landscape.

### 4.1. Factors influencing intraspecific variation

Limited evidence suggests that individual variations within populations, including social hierarchy and differences in morphology and behaviour, can influence seed dispersal patterns (Bartel & Orrock, 2021; Zwolak & Sih, 2020). The evidence on intraspecific variation in seed dispersal indicates contrasting results in the diversity of food items consumed by adults versus juveniles. There is limited previous information on intraspecific variation in the diversity of fruit items consumed by members of different age-sex categories. Juvenile howler monkeys *Alouatta pigra* fed on a lower diversity of fruits than adults (Benítez-Malvido et al., 2014). In contrast, we found that sub-adult LTMs consumed a greater diversity of fruits than adult male LTMs. While there was no difference in the richness of seeds dispersed by Japanese Macaques (Tsuji, Campos-Arceiz, et al., 2020), information on the diversity of fruits they consumed is absent.

Factors influencing intraspecific variation in fruit consumption have been attributed to greater experience in adult howler monkeys (Benitez-Malvido et al., 2016). This variation could also be influenced by dominance hierarchies, as observed in Japanese macaques (Tsuji, Campos-Arceiz, et al., 2020). Behavioural differences in social status and foraging patterns have been reported across different age-sex categories of LTMs (Dhawale et al., 2020; Singh et al., 2011). Adult male LTMs are dominant and display agonistic behaviour towards females and other males, indicating their dominance over females and subadults (Singh et al., 2011). This could compel the subadults to forage on a greater diversity of resources. Apart from the role of dominance in influencing intraspecific differences in the diversity of fruit items in LTM diets, these differences may result from variations in time spent foraging. Unlike previous studies, we found that adult males spent less time foraging and more time resting than subadults and females (Fig. S1). An animal that spends more time foraging and less time resting is likely to encounter more resources, which can explain the greater diversity of resource items in diets of subadults and females, an aspect that needs further investigation.

### 4.2. Implications of intraspecific variations on frugivory and seed dispersal

To determine seed dispersal effectiveness, we evaluated both quantitative (number of seeds dispersed) and qualitative measures (fruit-handling behaviour, movement patterns of LTMs and seed deposition sites). We could determine the number of seeds dispersed only for *Ficus* spp. While the low representation of medium- and large-seeded species like *Artocarpus heterophyllus* is expected since LTMs mostly spit seeds of these species, the low representation of *Lantana camara* seeds was surprising, given their extensive consumption of *Lantana*. The sieve size used for determining the diversity and number of seeds in LTM faeces followed established methods (Tsuji, Campos-Arceiz, et al., 2020) and was smaller than *Lantana camara* seeds. Macaques are known to grind seeds more than swallowing and defecating them intact (David et al., 2015; Stevenson, 2007). Therefore, it is likely that *Lantana* seeds are crushed, while the smaller *Ficus* seeds escape getting crushed.

Animals mainly disperse *Ficus* seeds in a clumped manner through their faeces (Zhou & Chen, 2010). There is no evidence indicating the negative impacts on germination or establishment due to clumped dispersal of *Ficus* seeds. Males showed a higher representation of *Ficus* in their diet compared to females and subadults (Fig. S2). Although females and subadults consumed a similar proportion of *Ficus* fruits, the number of *Ficus* seeds dispersed in the scats was lower in subadults than in females. This is likely due to physiological differences in fruit processing among the age-sex categories, which needs further investigation.

The quality of seed dispersal services is determined by fruit handling behaviour, dispersal distance and microhabitats where the seeds are dispersed (Schupp et al., 2010). All three age-sex categories had a low representation of unripe fruits. Therefore, they mainly disperse mature seeds of species whose seeds are not destroyed by them (e.g., seeds of *Artocarpus* are also predated). When macaques spit seeds, the dispersal distances are generally short (∼20 m), but defecated seeds may be dispersed up to 250 m from the parent tree (Tsuji & Su, 2018). Thus, macaques mainly disperse the seeds away from the parent tree, especially when they defecate and disperse. We did not find differences in movement patterns across the age-sex categories, suggesting that they disperse seeds at similar distances unless there are systematic differences in seed retention times, which is currently unknown. The most suitable microsites for *Ficus* seed germination and seedling establishment are tree trunks and branches (Laman, 1995; Shanahan, 2016). We found weak evidence that male macaques tend to defecate on trees, thereby they are more likely to disperse *Ficus* seeds in favourable microsites.

### 4.3. Implications on dispersal of non-native species

In disturbed landscapes, primates feed on fruits of exotic invasive species, contributing more than 50% of their diet (Canale et al., 2016; Wimberger et al., 2017) and are major dispersers of these alien species (Cordeiro et al., 2004; Dew & Wright, 1998; Silva et al., 2018). While there is substantial evidence of frugivory and seed dispersal of alien invasive species by primates, there is limited literature on intraspecific variation. Given the long history of human use and associated forest degradation in our study area, there is a significant presence of non-native species, including alien invasives like *Coffea* and *Lantana*, which are among the most problematic invasives in the area (Joshi et al., 2009, 2015). Interestingly, unlike females and subadults, males consumed a greater diversity of non-native fruits than native fruits, highlighting their role in dispersal of invasive plants, especially coffee. The reason for intraspecific variation in consumption of *Lantana*, which was consumed more by females and subadults, is unclear.

*Coffea* berries are more energy-rich, with higher lipid and protein content than *Lantana* berries (Embaby & Mokhtar, 2011; Osorio Pérez et al., 2023) differences. Like most primates which consumed the pulp of the exotics, dropping seeds under the mother tree (Canale et al., 2016; Cordeiro et al., 2004), LTMs spat *Coffea* seeds undamaged mainly under the mother plant (pers. obs.). However, the tiny *Lantana* seeds were probably destroyed by them. While there are reports of *Lantana* fruit consumption by LTMs (Singh, Kumara, et al., 2002), no confirmed reports exist on the effective dispersal of *Lantana* seeds by LTMs. We found only one report of *Lantana* dispersal by *Papio anubis* (Simon et al., 2016). Given the differences in relative abundances of males, females, and subadults in the troops, the overall number of seeds dispersed by these different age-sex categories is likely to differ.

The rainforest fragment where the study was conducted is highly degraded, yet supports among the highest recorded densities of LTMs. The densities of native rainforest trees were significantly lower in the focal study area than in Protected Areas, but the density of non-native trees was substantially higher (Joshi et al., 2009, 2015). Thus, given the low density of native trees in this fragment, alien species are important food resource for the primates. While previous studies have suggested that provisioning may contribute to higher densities of LTMs in this fragment (Dhawale et al., 2020), the hyperabundance of *Coffea* and *Lantana* fruits can also contribute to their relatively higher densities. This aspect must be investigated by comparing native and non-native fruit availability between rainforests and degraded fragments with alien species over space and time to assess their role in supporting primate populations.

This study adds to the growing literature on intraspecific variation in seed dispersal highlighting the role of age and sex on the quantitative and qualitative contributions to seed dispersal of native and non-native plants. While our sampling effort exceeds that of Tsuji et al. (2020), it is restricted to a single season. Inter-annual variation in fruiting in the temperate forests of Japan was found to influence the intraspecific variation in seed dispersal. Given the interannual variation in fruit availability in tropical forests (Mendoza et al., 2018), similar studies need to be conducted to determine intraspecific variation in seed dispersal across years.

Moreover, our study was conducted in a disturbed rainforest fragment (due to the logistical challenges of conducting focal observations on primates, as only this population is habituated to human observers in the landscape). A similar study in more intact forests with a lower prevalence of invasive species can provide vital insights into the influence of broader contexts on intraspecific variation in seed dispersal.

## AUTHOR CONTRIBUTIONS

RN, BK and HNK conceived ideas and designed the study; BK collected data; BK and RN carried out data analysis with inputs from HNK; BK and RN led the writing of the manuscript with inputs from HNK.

## ACKNOWLEDGEMENTS

This study was part of the Master’s dissertation at the Wildlife Institute of India. We thank the Tamil Nadu Forest Department for granting the permits to carry out the study. BK thanks Navendu for his support and guidance. We are grateful to Ashni Dhawale for her valuable feedback. BK thanks Nature Conservation Foundation for support during fieldwork and manuscript writing. We thank the Scientific and Engineering Research Board (Anusandhan National Research Foundation; Grant no: SRG/2021/001523), Rohini Nilekani Philanthropies for partially funding the fieldwork. We thank Kim McConkey for feedback on the thesis and H. S. Sushma for fruitful discussions. BK thanks Divya and Shankar Raman for their valuable comments and support during the fieldwork. BK thanks Srinivasan, Vishnu, and Satish for helping with logistics and fieldwork. BK thanks Madhavan for identifying *Ficus* species. BK thanks Monika, Qamar Qureshi, Joel, Mohit Mudliar, Aditya Satish, Arnab and Abhishek for useful discussions and help.

## DATA AVAILABILITY

Data will be uploaded on Zonedo repository upon acceptance of the manuscript.

---

**Figure S1.**
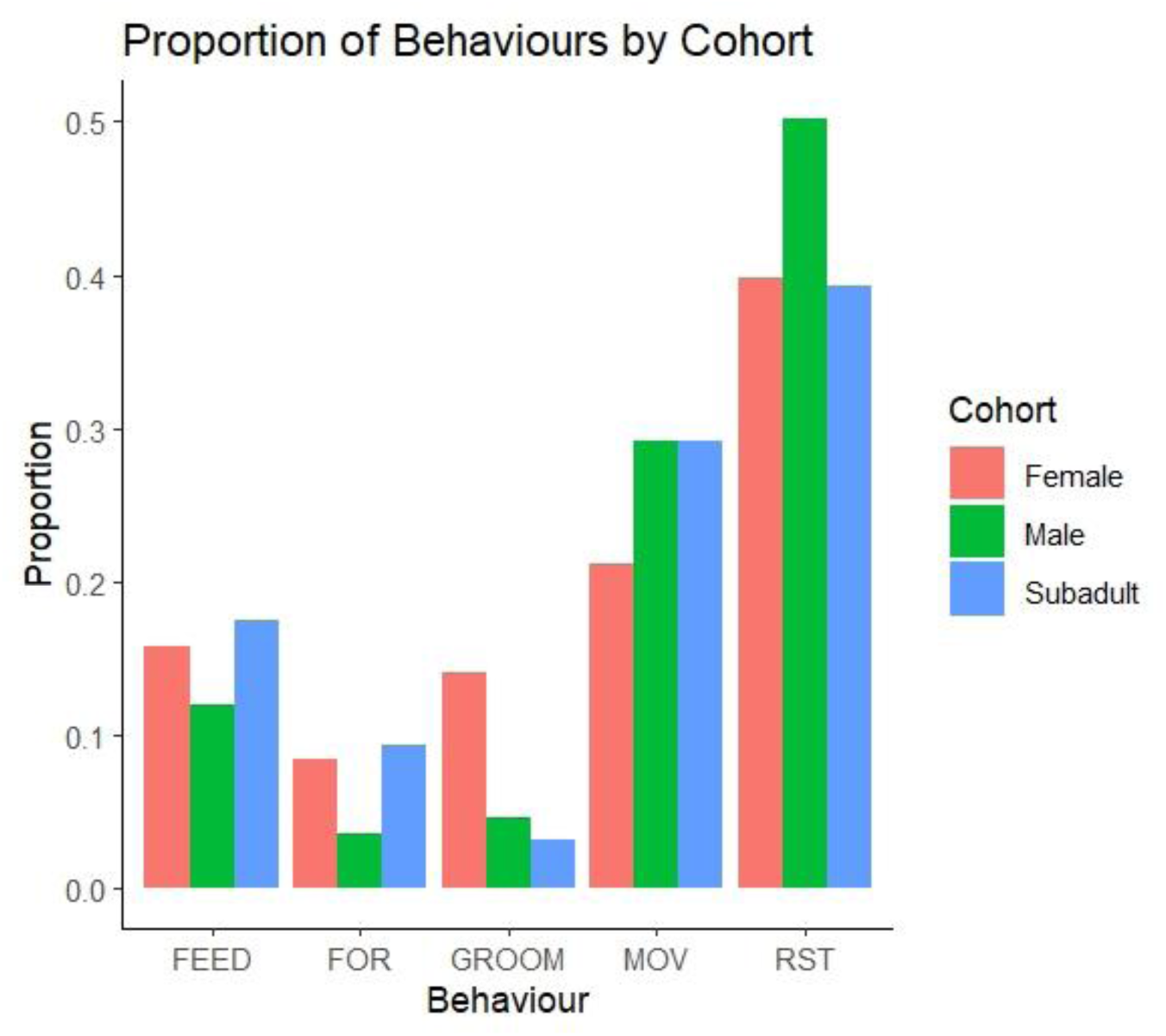
Bar plot showing the time spent on different behaviours by the three age-sex categories of lion-tailed macaques. Activities are abbreviated as FEED - Feeding, FOR - Foraging, GROOM - Grooming, MOV - Moving, RST - Resting. Foraging behaviour includes searching for food items in litter, bark, leaves or other substrate. We ran beta regression models to examine if there was a significant difference in time spent on feeding, foraging and resting behaviours across the age-sex categories. We found males were foraging less and resting more than females and subadults. There was no difference in time spent on feeding by the different age-sex categories.

**Figure S2.**
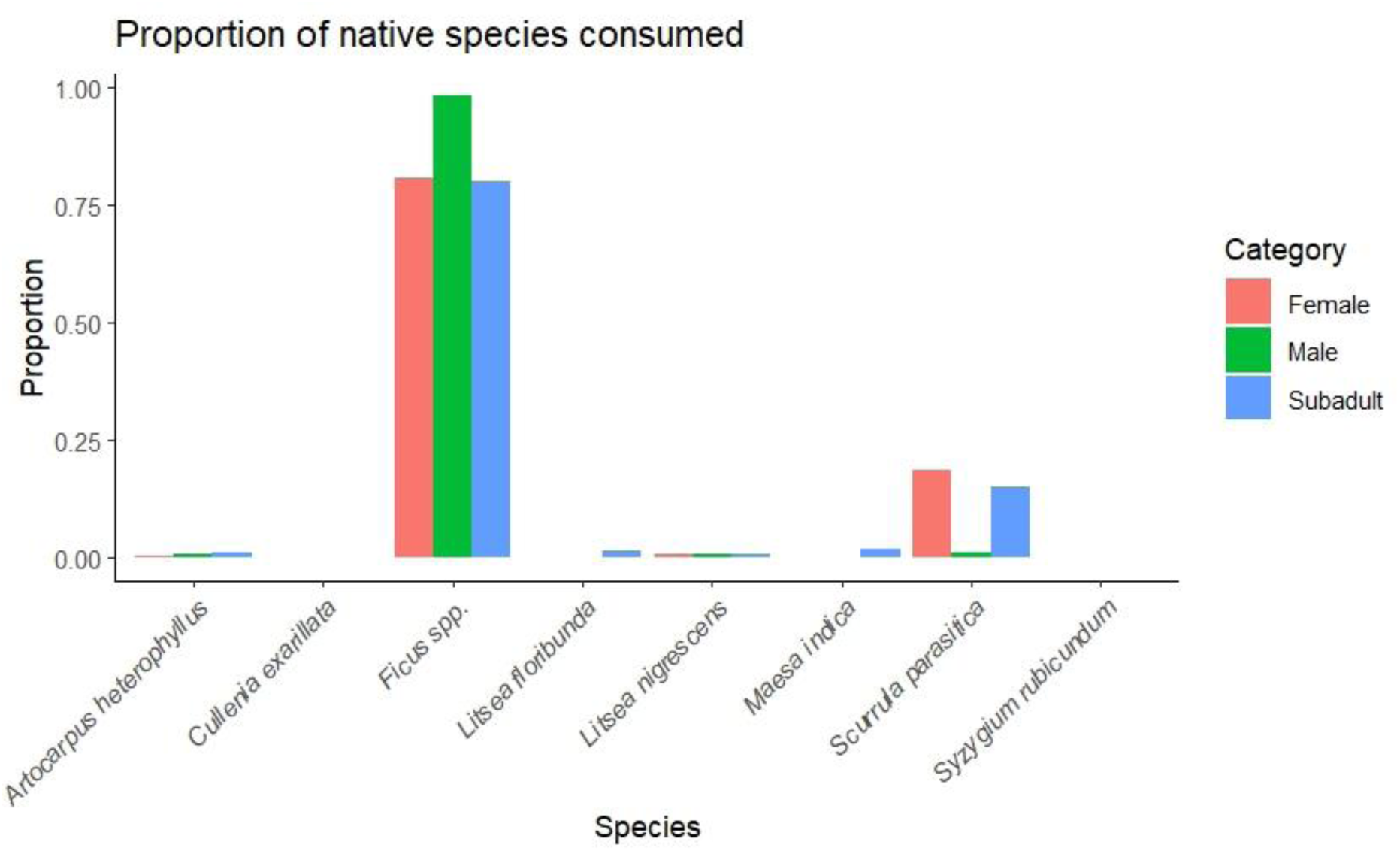
Bar plot showing the relative proportion of native fruit species consumed by different age-sex categories of lion-tailed macaques. The nine species of *Ficus* recorded in the diet were pooled as *Ficus* spp. Chi-square test was performed to examine the differences in the proportion of native species consumed across age-sex categories. Males consume higher proportion of *Ficus* spp. than females and subadults (*ꭓ^2^* = 1156.2, df = 14, *p* < 0.001).

**Table S1.** Age-sex classification of lion-tailed macaques (adapted from Singh et al., 2002)

| Age-sex class | Age range |
| --- | --- |
| Adult male | > 8 years |
| Subadult male | 4–8 years |
| Adult female | > 6 years |
| Subadult female | 4–6 years |
| Juvenile | 1–4 years |
| Infant | <1 year |

**Table S2.**
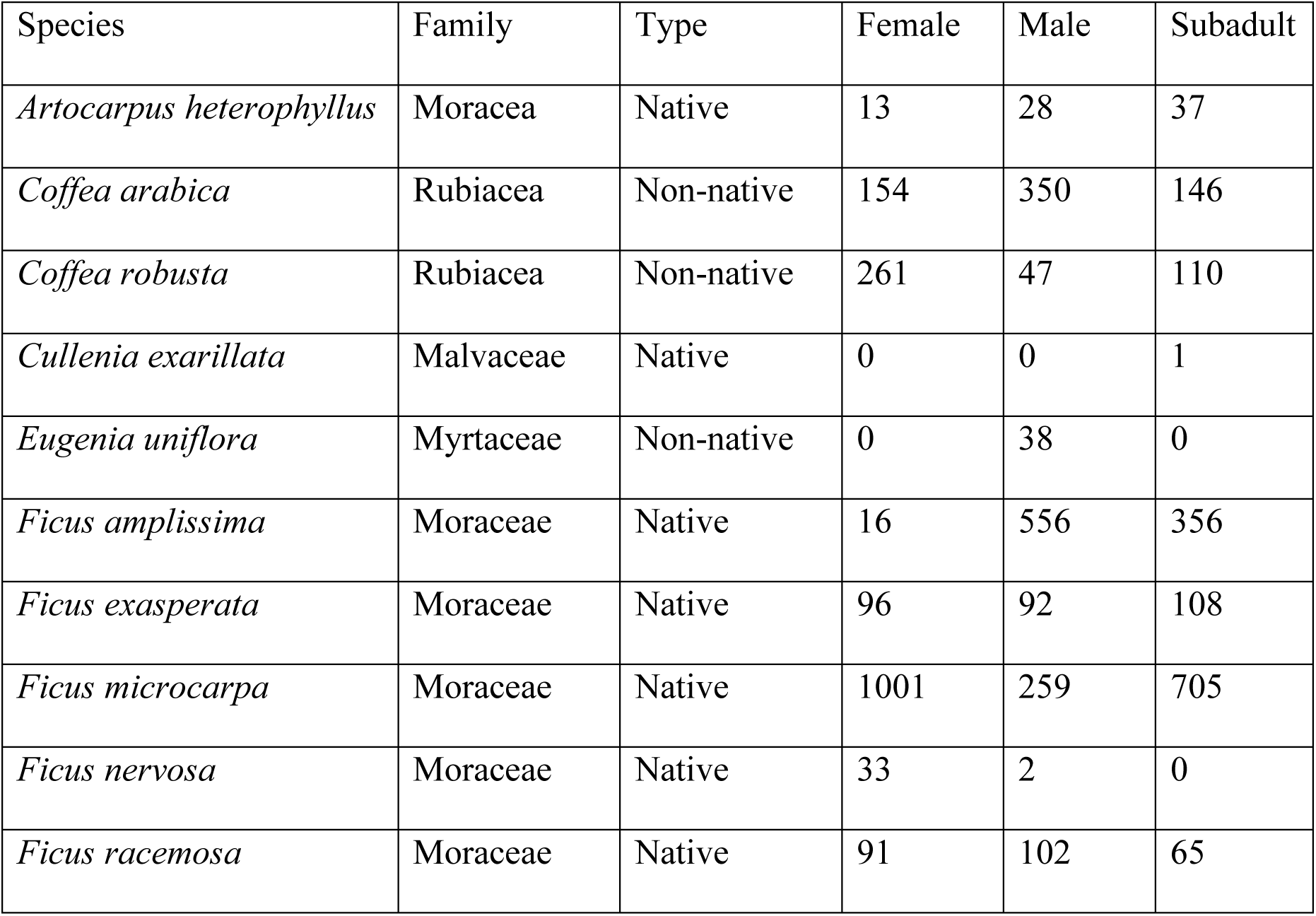
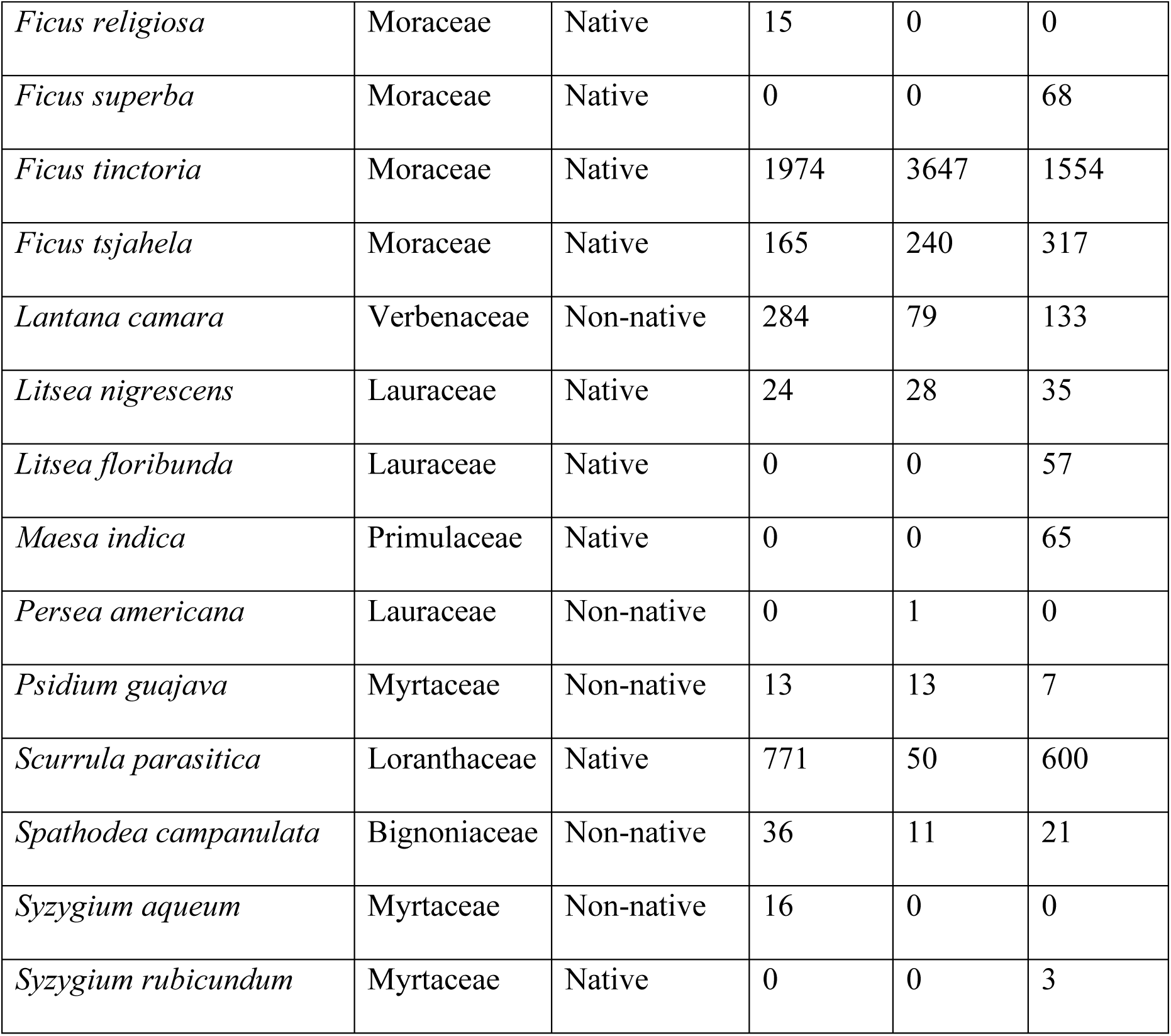
Number of fruits of different plant species that we observed the different age-sex categories of Lion-tailed macaques feeding on during the study period.

| Species | Family | Type | Female | Male | Subadult |
| --- | --- | --- | --- | --- | --- |
| *Artocarpus heterophyllus* | Moraceae | Native | 13 | 28 | 37 |
| *Coffea arabica* | Rubiaceae | Non-native | 154 | 350 | 146 |
| *Coffea robusta* | Rubiaceae | Non-native | 261 | 47 | 110 |
| *Cullenia exarillata* | Malvaceae | Native | 0 | 0 | 1 |
| *Eugenia uniflora* | Myrtaceae | Non-native | 0 | 38 | 0 |
| *Ficus amplissima* | Moraceae | Native | 16 | 556 | 356 |
| *Ficus exasperata* | Moraceae | Native | 96 | 92 | 108 |
| *Ficus microcarpa* | Moraceae | Native | 1001 | 259 | 705 |
| *Ficus nervosa* | Moraceae | Native | 33 | 2 | 0 |
| *Ficus racemosa* | Moraceae | Native | 91 | 102 | 65 |
| *Ficus religiosa* | Moraceae | Native | 15 | 0 | 0 |
| *Ficus superba* | Moraceae | Native | 0 | 0 | 68 |
| *Ficus tinctoria* | Moraceae | Native | 1974 | 3647 | 1554 |
| *Ficus tsihela* | Moraceae | Native | 165 | 240 | 317 |
| *Lantana camara* | Verbenaceae | Non-native | 284 | 79 | 133 |
| *Litsea nigrascens* | Lauraceae | Native | 24 | 28 | 35 |
| *Litsea floribunda* | Lauraceae | Native | 0 | 0 | 57 |
| *Maesa indica* | Primulaceae | Native | 0 | 0 | 65 |
| *Persea americana* | Lauraceae | Non-native | 0 | 1 | 0 |
| *Psidium guajava* | Myrtaceae | Non-native | 13 | 13 | 7 |
| *Scurrula parasitica* | Loranthaceae | Native | 771 | 50 | 600 |
| *Spathodea campanulata* | Bignoniaceae | Non-native | 36 | 11 | 21 |
| *Syzygium aqueum* | Myrtaceae | Non-native | 16 | 0 | 0 |
| *Syzygium rubicundum* | Myrtaceae | Native | 0 | 0 | 3 |

**Table S3.** Results of General Linear Model (Gaussian error structure) examining differences in the distance travelled across age-sex categories.

| **Predictor** | **Estimate** | **SE** | **t** | **p** |
| --- | --- | --- | --- | --- |
| Intercept: Female | -0.2766 | 0.04223 | -6.551 | **<0.001** |
| Male | 0.00764 | 0.05761 | 0.133 | 0.895 |
| Subadult | 0.06607 | 0.05626 | 1.174 | 0.243 |

**Table S4.** Results of the Generalised Linear Model (negative binomial error structure) examining differences in the number of *Ficus* seeds dispersed across age-sex categories.

| Predictor | Estimate | SE | t | p |
| --- | --- | --- | --- | --- |
| Intercept: Female | 8.064 | 0.241 | 33.403 | **<0.001** |
| Male | -0.074 | 0.346 | -0.214 | 0.830 |
| Subadult | -0.766 | 0.379 | -2.021 | **0.043** |

